# Structured Neural Variability from Repeated Naturalistic Video Watching Experiences

**DOI:** 10.1101/2025.11.07.687258

**Authors:** Menghan Yang, Pin-hao A. Chen, Amanda M. Brandt, Sushmita Sadhukha, Max Farrens, Eshin Jolly, Luke J. Chang

## Abstract

The brain continuously integrates perception and interpretation processes to navigate dynamic environments. It is often assumed that these processes are invariant to repeated experiences. Here we investigated how neural responses to naturalistic stimuli evolve across multiple viewings using functional magnetic resonance imaging. Twenty participants watched 24 short videos across five separate scanning sessions. Between-subject correlations (BSC) decreased progressively across all brain regions with repeated viewing, indicating increasing individual variability. Within-subject correlations (WSC) consistently exceeded BSC, demonstrating that individuals are more similar to themselves than to others. However, the structure of within-subject variability differed fundamentally between sensory and interpretive regions. Early visual cortex exhibited strong repetition effects, with brain activity becoming increasingly dissimilar with each successive viewing, accompanied by reduced coupling to low-level visual features and decreased anti-correlation with default mode regions. In contrast, ventromedial prefrontal cortex (vmPFC) showed temporal proximity effects, where sessions closer in time exhibited more similar activity patterns regardless of viewing number. This temporal structure in vmPFC aligned with fluctuations in participants’ video preferences across sessions. These findings suggest that repeated experiences are never processed identically: sensory systems adapt efficiently to familiar input while interpretive systems continuously reshape meaning based on current internal states.

## Introduction

Our brains are remarkably adept at creating and executing complex plans in dynamic, uncertain environments, striving to maximize survival while minimizing metabolic costs. This intricate process begins with **perception**, where exogenous signals from the external world are transduced into neural signals within unimodal primary sensory cortices (Margulies et al., 2016; Mesulam, 1998; Zhang et al., 2024). These perceptual signals are then subject to **interpretation**, in which their meaning is evaluated in the context of endogenous signals representing internal states and goals (FeldmanHall & Chang, 2018; O’Reilly et al., 2014), primarily within transmodal ventromedial prefrontal cortex (vmPFC; (Ashar et al., 2017; Morecraft et al., 1992; Roy et al., 2012; Yeshurun et al., 2021)). The dynamic interplay between these perception and interpretation systems ultimately shapes how we navigate and respond to our environment.

Understanding how these perception and interpretation systems interact in real-world contexts requires methodological approaches that capture their complex interactions. Naturalistic passive movie watching paradigms have gained popularity for their ability to simultaneously probe multiple interacting cognitive processes, complementing traditional approaches that focus on isolating specific cognitive processes (Finn et al., 2020; Finn & Bandettini, 2021; Sonkusare et al., 2019). Research employing these paradigms has consistently demonstrated that perceptual activity in unimodal sensory cortices is largely shared across individuals (Hasson et al., 2004; Lu et al., 2016; Mukamel et al., 2005), as measured by inter-subject correlation analysis (ISC; (Nastase et al., 2019)). In contrast, activity in transmodal regions like the vmPFC appear to be highly idiosyncratic across individuals consistent with its hypothesized role in interpreting the meaning of this information (Nguyen et al., 2019) with respect to past experiences, current homeostatic states, and future goals (L. J. Chang et al., 2021; Xie et al., 2021).

Beyond distinguishing shared perceptual responses from idiosyncratic interpretive processes, an open question is how repeated experiences shape these dynamics. Although movie-watching paradigms are widely used to study neural development (Richardson et al., 2018; Vanderwal et al., 2019), clinical symptoms (Byrge et al., 2015; Eickhoff et al., 2020; Kirk et al., 2024; Lahnakoski et al., 2022), and individual differences (Finn & Bandettini, 2021; Vanderwal et al., 2017), the field has yet to systematically investigate how repeated viewing affects neural responses. In contrast to traditional tasks, where test-retest reliability (Cronbach, 1951) is routinely assessed (Bennett & Miller, 2013; Demidenko et al., 2024; Friedman et al., 2008; Han et al., 2022; Korucuoglu et al., 2020), the reliability of responses to repeated movie exposure remains largely unexamined—raising the open question of how repeated experiences reshape perceptual and interpretive processes. The limited extant work suggests that temporal dynamics of voxel activity in unimodal visual and auditory cortex are largely preserved with repeated viewing (Andric et al., 2016; Golland et al., 2007; Hasson et al., 2010; Wang et al., 2017; Wilf et al., 2017) as well as the structure of individual differences in these responses (Gao et al., 2020). Transmodal regions, however, exhibit greater temporal instability, with patterns anticipating future events shifting up to 15 seconds prior to event boundaries (Lee et al., 2021).

How, then, might brain processes within the same individual change as a function of repeated viewing? Research on perception using traditional task-based experimental designs demonstrates that repeated presentations of the same stimulus do not always elicit identical neural responses. Repetition suppression, for example, is a phenomenon in which immediately repeated stimulus presentations lead to temporarily reduced neural activity in sensory cortices (Grill-Spector et al., 2006), potentially reflecting the influence of expectations on perception (Summerfield et al., 2008). This phenomenon can be understood through the lens of predictive coding, which posits that the brain continuously generates predictions about incoming sensory inputs and uses prediction errors (the difference between predicted and observed input) to refine its internal model of the external world (Auksztulewicz & Friston, 2016). This process aligns with the free energy principle, which suggests that minimizing surprise is a metabolically efficient way to optimize neural computations and behavior (Friston, 2010). Thus, from this perspective, repeated movie viewing might lead to *increased* changes in activity patterns as the brain detects new information and updates its predictions to minimize uncertainty.

In contrast to perception, *interpretation* involves generating evaluative appraisals of sensory information based on past experiences, current needs, and future goals. If these endogenous signals remain stable across repeated experiences, the interpretation of sensory information might also remain consistent. However, changes in these internal states are likely to alter the interpretation of sensory input. For example, the smell of food is likely to be interpreted differently when an agent is hungry versus satiated (Panksepp, 2004). Research on value-based decision-making has consistently implicated the vmPFC in generating the subjective value of a stimulus (Gottfried et al., 2003; Kringelbach et al., 2003; O’Doherty et al., 2002; Padoa-Schioppa & Assad, 2006; Plassmann et al., 2007) by integrating multiple stimulus features (S. Suzuki et al., 2017). However, this appraised value is sensitive to changes in homeostatic states (Howard & Kahnt, 2017; Tomova et al., 2020) and broader goals (Hare et al., 2008; Plassmann et al., 2010). Because memories, goals, and homeostatic states fluctuate across different timescales, interpretations of the same sensory experience may drift depending on the temporal context (Manning, 2021; Manning et al., 2015; Yeshurun et al., 2017). Watching the same movie in the same week, for instance, is likely to elicit more similar interpretations than watching it during childhood and again in adulthood.

In the present study, we aimed to characterize how brain activity within individuals is impacted by repeated exposure to naturalistic stimuli using a passive movie-watching paradigm. Participants (N=20) underwent functional magnetic resonance imaging (fMRI) while watching 24 short videos selected to elicit a variety of affective experiences, across five separate scanning sessions spanning three hours. We specifically investigated how activity in unimodal and transmodal cortices changed with repeated experiences, both within and across individuals.

## Results

### Assessing neural stability across repeated video-watching experiences

Our initial goal was to establish a baseline measure of the similarity in brain dynamics across participants while watching the same movie. To achieve this, we computed temporal pairwise between-subject correlations (BSC) for 50 distinct regions defined by a meta-analytic parcellation (de la Vega et al., 2016). This analysis was performed separately for each of the 24 videos and each of the 5 viewing sessions. Hypothesis tests, performed over subjects, revealed patterns consistent with previous research (L. J. Chang et al., 2021; Hasson et al., 2004; Lerner et al., 2011; Simony et al., 2016). Specifically, during the first viewing session, we observed higher BSCs in primary sensory cortices and lower correlations in heteromodal and transmodal cortices (Fig 1A, top). However, the average temporal BSC across all viewing sessions appeared to decrease across the entire brain (Fig 1A, middle). This decrease was statistically significant in both V1 (*t*_(23)_ = -15.354, *p* < 0.001) and vmPFC (*t*_(23)_ = -5.509, *p* < 0.001, Figure 1B blue and green bars), suggesting increased individual variation with each successive viewing (Figure 1C; V1: linear contrast *t*_(23)_ = -19.002, *p* < 0.001) ; vmPFC: linear contrast *t*_(23)_ = -6.430, *p* < 0.001).

**Figure 1:**
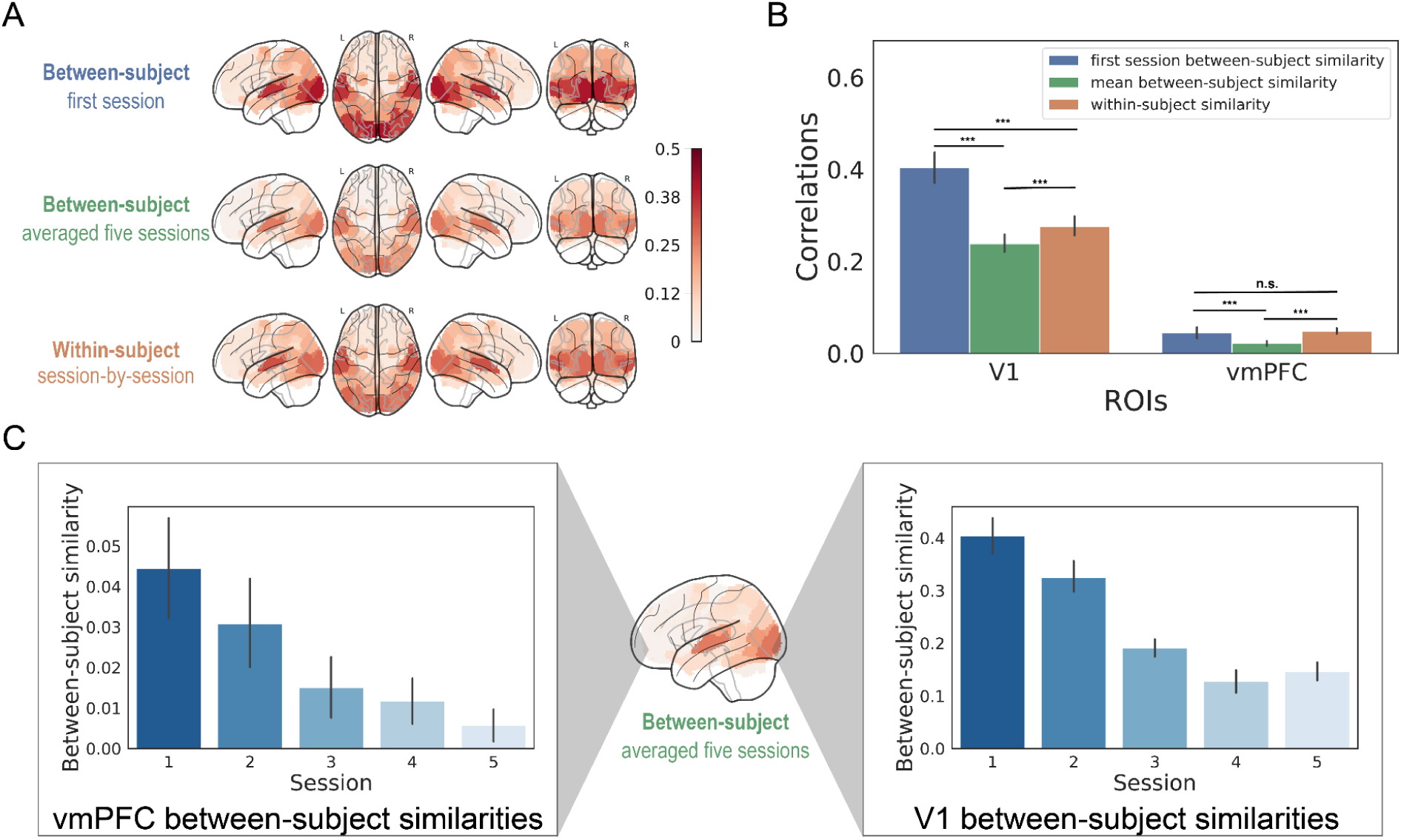
Compare between-subject correlations (BSC) and within-subject correlations (WSC) across repeated video-watching. (A) Whole-brain maps of BSC during the first session (top), average BSC across all five sessions (middle), and WSC across repeated sessions (bottom). The overall spatial patterns of mean BSC and WSC across parcels were highly similar (also see Figure S1). (B) Unimodal and heteromodal ROI comparison for WSC and BSC effects. ROI analyses revealed distinct patterns across regions. In V1, first-session BSC was highest, WSC was intermediate, and mean BSC across sessions was lowest (*p*s < 0.001 for all pairwise comparisons). In vmPFC, BSC and WSC values were overall lower than in V1, but WSC was still significantly greater than mean BSC (*p* < 0.001), while not differing from first-session BSC. These results indicate that people are more similar to themselves than to others when viewing the same stimuli. (C) Session-by-session analyses showed that BSC declined across repeated viewings in both V1 and vmPFC, indicating increasing individual variability across sessions. Error bars indicate 95% confidence intervals.

Next, we examined the average within-subject correlation (WSC) across the repeated viewing sessions. For each brain region, we calculated the temporal WSC for each pair of viewing sessions separately for each video, and then averaged these correlations across videos and participants. Interestingly, the broad spatial pattern of WSC across parcels closely mirrored the average BSC (Figure S1C, mean *r* = 0.966, sd=0.010, *t*_(23)_ = 455.696, *p* < 0.001). This suggests that regions exhibiting similar temporal dynamics across participants also show consistent activity patterns within the same participant across repeated viewings. Unimodal visual and auditory cortices demonstrated the highest within-subject reliability (Andric et al., 2016; Golland et al., 2007). Furthermore, we observed a significantly overall higher degree of synchronization within the same participant compared to between subjects averaged across the five viewing sessions (Figure 1C) in both V1 (*t*_(23)_ = 12.152, *p* < 0.001) and vmPFC (*t*_(23)_ = 8.808, *p* < 0.001). This finding aligns with the notion that people are more similar to themselves than to others when viewing the same stimuli.

### Decomposing the structure of within-individual variability

We next turned our attention to characterizing changes in brain activity within individuals across repeated viewings of the same movie. We computed the average within-subject temporal similarity of activity within brain regions across viewing sessions to create within-subject temporal representational similarity matrices and identified distinct patterns of change in unimodal (V1) and transmodal (vmPFC) cortices (Figure 2A). In V1, activity patterns appeared to change as a function of the number of times a video was viewed. Early viewing sessions exhibited greater similarity to each other and less similarity to later sessions, consistent with the “Anna Karenina” pattern (Finn et al., 2020). In contrast, vmPFC activity exhibited a pattern more consistent with temporal autocorrelation, where sessions closer in time showed more similar temporal dynamics, irrespective of the viewing number.

**Figure 2:**
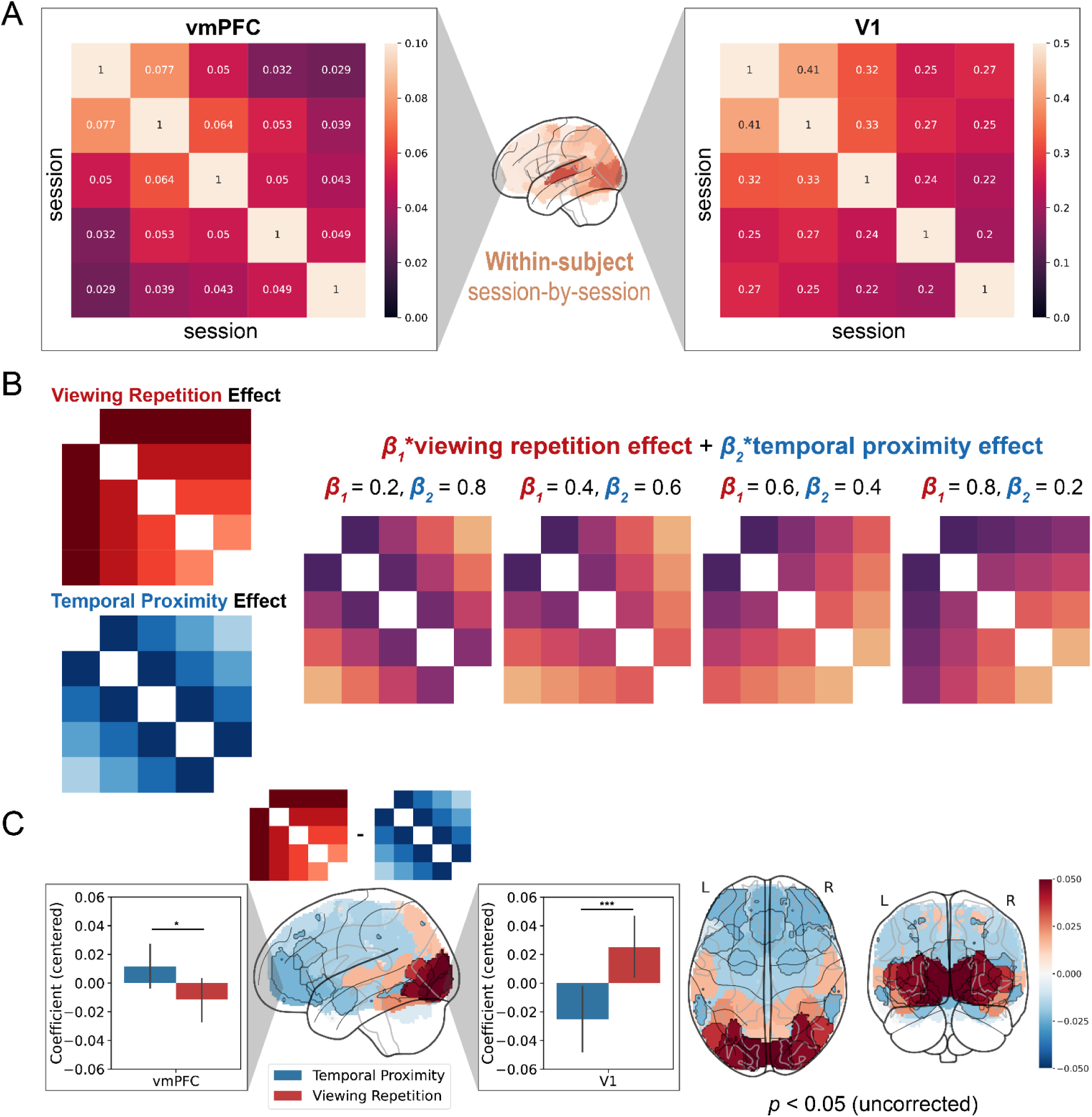
Decomposing the structure of within-subject variability across repeated video-watching. (A) Average session-by-session within-subject representational similarity matrices for V1 and vmPFC. V1 showed higher similarity among early sessions and divergence from later sessions (repetition effect), whereas vmPFC showed stronger similarity between temporally adjacent sessions (temporal proximity effect). (B) Prototypical matrices representing Viewing Repetition and Temporal Proximity effects, and simulated similarity matrices generated by their linear combinations. (C) We applied distance regression and the contrasts between repetition and proximity effects revealed stronger repetition effects in V1 (*t* = –3.589, *p* < 0.001, right bar plot) and stronger temporal proximity effects in vmPFC (*t* = 2.112, *p* = 0.035, left bar plot). Whole-brain analysis across 50 parcels showed similar dissociations, with only V1 and V2 surviving FDR correction (*q* < 0.05). Error bars represent bootstrapped 95% confidence intervals.

To formally quantify these observations, we employed a distance regression approach (Parkinson et al., 2017; Xie et al., 2021) to decompose the observed within-subject representational similarity matrix for each video from two prototypical matrices: temporal proximity and viewing repetition. The temporal proximity effect was modeled using a Toeplitz matrix, with each descending diagonal monotonically decreasing to capture autocorrelation across viewing sessions. The viewing repetition effect was modeled by a matrix that monotonically decreased as a function of distance from the first viewing session. Figure 2B illustrates simulated examples of representational similarity matrices that could be generated by linear combinations of these two prototypical matrices. We estimated *β* weights for each of these matrices for each participant and each video by finding the optimal linear combination that best reconstructed the observed representational similarity matrix via least squares minimization.

We then used a mixed effects regression model to test whether different regions exhibited distinct patterns of temporal proximity and viewing repetition effects, including random intercepts for videos and participants. While both V1 and vmPFC showed variations influenced by repeated viewing times and session proximity (Figure S1 and S2), the impact of these two effects differed significantly between the regions (*F*_ROI*effect_ = 16.160, *p* < 0.001). Specifically, V1 exhibited a stronger repetition effect (*t* = -3.589, *p* < 0.001), indicating that responses became more variable with each successive viewing. Conversely, vmPFC showed a stronger temporal context effect (*t* = 2.112, *p* = 0.035), suggesting that increased temporal proximity between two viewing sessions might evoke more similar endogenous signals and overall interpretations of the narrative.

### Sensitivity Analyses

To ensure the robustness of these findings, we conducted several additional analyses. First, we relaxed the assumption of equally spaced viewing repetitions in our distance regression model. Due to scheduling variability, the actual time between scanning sessions varied across participants (mean interval = 11.0 days, SD = 3.57, Figure S4A). We repeated the same analyses using the actual time intervals measured in days (Figure S4B) and found similar results: temporal repetition effects were stronger in unimodal cortex, while temporal context effects were stronger in transmodal cortex (Figure S4C). Second, we investigated whether similar effects could be observed in spatial patterns, extending beyond our primary focus on temporal dynamics. We conducted the same analyses using spatiotemporal dynamics, creating time-by-time similarity matrices to capture how spatial patterns vary within each participant over time (L. J. Chang et al., 2021). We then averaged the time-varying similarity over all time points within a movie, and then across movies. We successfully replicated our neural stability analyses (Figure S5) as well as our within-subject decomposition analyses (Figure S6) using this spatiotemporal approach, demonstrating the robustness of our findings to the inclusion of spatial pattern information.

### Bridging within-subject variability to cognitive processes

Finally, we sought to establish a more direct link between our neural findings and underlying cognitive processes. First, we hypothesized that the decreased WSC observed in unimodal areas like V1 might reflect reduced perceptual engagement with repeated viewings, possibly due to habituation or a shift in attentional focus which would manifest as a change in a relationship to a visual property of the stimulus (Grill-Spector et al., 2006), To test this, we computed the average pixel intensity for each video frame and downsampled this luminance timeseries to 0.5Hz to match the fMRI data’s sampling rate. We then used a GLM to estimate how well each voxel in the brain could be explained by one of the most basic properties of each image, image brightness, separately for each video and each session. We used a linear contrast to test for effects that monotonically decreased over viewing sessions with an FDR threshold of q < 0.05 (Figure 3A). Even though image luminance is identical across all viewing sessions, we found a significant decrease in V1 activation (*β* = -0.834, *t* = -64.379, *p* < 0.001), indicating a reduced association with visual features over successive viewing sessions. Voxelwise results showed this decrease extended to adjacent early visual areas including the occipital pole, lateral occipital cortex, and occipital fusiform gyrus. Interestingly, we found that the luminance effects in voxels in vmPFC (*β* = 0.047, *t* = 10.326, *p* < 0.001) increased from being initially anti-correlated to not being correlated with activity in these regions by the final viewing sessions. We also observed positive shifts in other heteromodal regions, including the angular gyrus, supramarginal gyrus, precuneus, and frontal pole. Covarying activity in these regions are often referred to as “default mode” activity and are associated with internal thought such as mind wandering (Christoff et al., 2016; Mason et al., 2007) that are anti-correlated with task-positive networks (Fox et al., 2005; Gusnard et al., 2001). We speculate that these findings may indicate a shift from external visual processing to internal or endogenous processing with increased stimulus familiarity.

**Figure 3:**
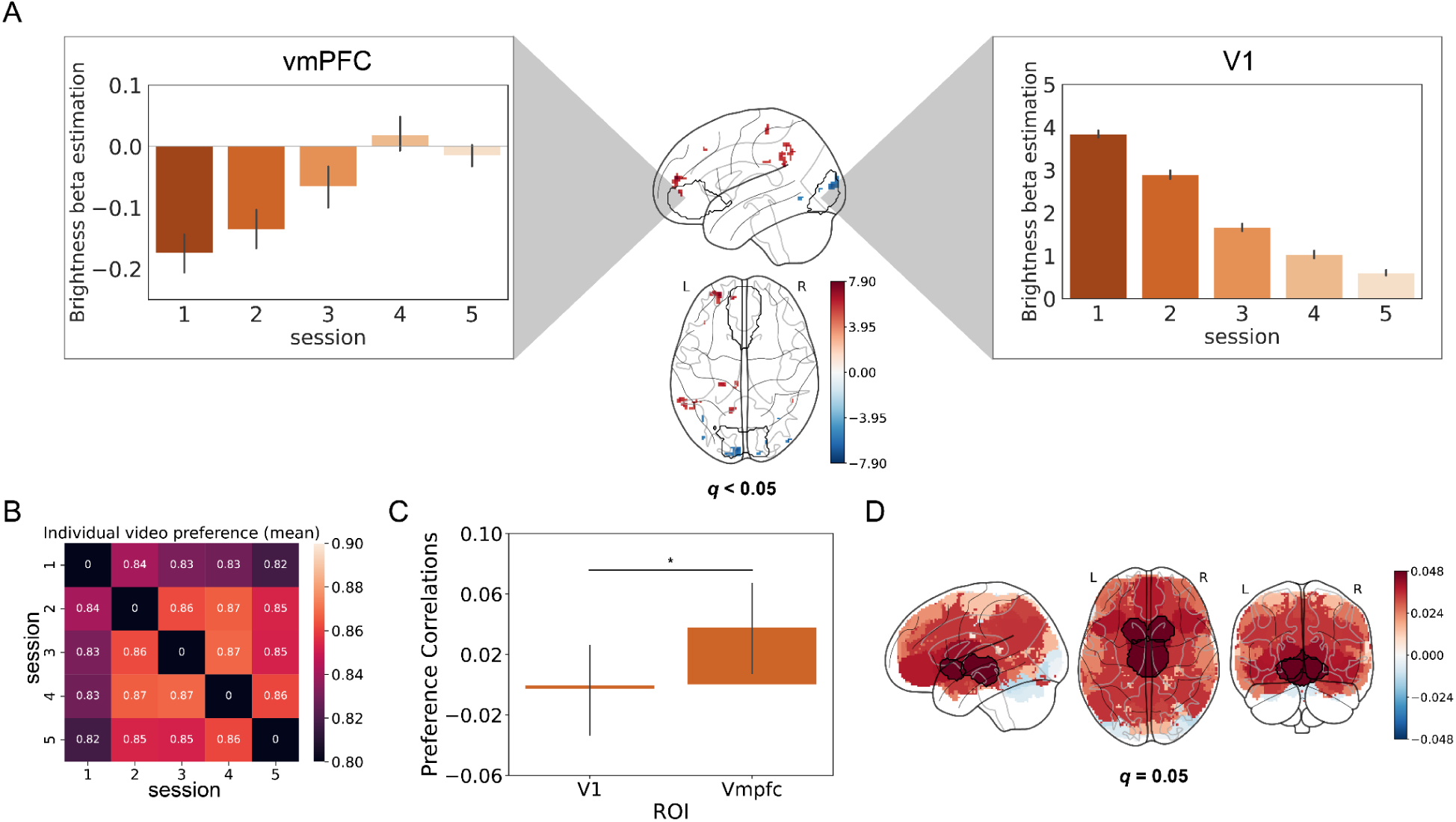
Variations in visual activation and individual video preferences across sessions and their neural representations. (A) We estimated whole-brain voxelwise activation to video luminance and tested monotonic linear changes across sessions. Significant voxels were identified after using FDR correction (*q* < 0.05). Parcellation results also showed increasing activation in vmPFC and decreasing activation in V1. (B) Matrix of individual video preferences (averaged across videos and participants). (C) Individual video preferences correlated more strongly with vmPFC than V1. (D) Correlations between individual video preferences and brain parcels. Nucleus accumbens and thalamus survived FDR multiple comparisons correction (*q* = 0.05).

Second, we aimed to connect the temporal context effects observed in transmodal cortex to behavioral measures of endogenous processing. While we lacked direct access to participants’ internal appraisals of each video, we assessed their preferences using a revealed preference task. After each scanning session, participants were presented with screenshots from two videos and asked to select their preferred video. We created a video-by-video revealed preference matrix for each session and computed how many times a participant preferred one video over all others to generate an ordinal ranking metric. Using representational similarity analysis (RSA), we mapped session-by-session ranked preferences (Figure 3B) to each brain parcel’s session-by-session temporal similarity matrix. We found that the vmPFC activity (mean *r* = 0.038, *p* = 0.005) was more reflective of changes in participants’ preferences compared to V1 (mean *r* = -0.003, *p* = 0.565; *t*_(19)_ = 2.035, *p* = 0.028, Figure 3C). A whole brain analysis revealed that subcortical areas, including the nucleus accumbens (Nacc) and thalamus, were the only regions to survive FDR multiple comparisons correction (Figure 3D).

## Discussion

The primary goal of this study was to characterize how the brain processes repeated sensory experiences. A central assumption in cognitive neuroscience is that individuals will generate similar internal experiences and neural responses when exposed to the same stimuli. We elicited a variety of affective experiences using 24 short videos viewed over 5 separate 3-hour scanning sessions. Overall, we observed that the reliability of brain activity across all videos followed the expected gradient from high synchronization in primary unimodal sensory cortices to greater idiosyncrasy in transmodal regions (L. J. Chang et al., 2021; Hasson et al., 2004; Nguyen et al., 2019). Furthermore, within-subject correlations consistently exceeded between-subject correlations across all regions and sessions (Andric et al., 2016; Gao et al., 2020; Golland et al., 2007; Wang et al., 2017). However, this gradient became progressively attenuated with repeated viewing, as both sensory and association cortex exhibited increased individual variability, suggesting that the entire cortical hierarchy adapts as participants become familiar with the stimuli. These results challenge the widely held assumption that experimental stimuli will consistently elicit the same neural and psychological processes.

Beyond demonstrating that repeated viewing leads to greater variability in neural dynamics, we also looked for systematic patterns in how activity within brain regions changed. Any structured patterns of change that are consistent across stimuli and participants could reveal insights into how regions process information elicited by the experimental stimuli. Our decomposition analysis revealed fundamentally different patterns of change in sensory versus transmodal cortices. Primary visual cortex (V1) exhibited a strong repetition effect, with neural responses becoming increasingly dissimilar with each successive viewing. This pattern aligns with established principles of predictive coding (Grill-Spector et al., 2006; Summerfield et al., 2008), suggesting that sensory systems optimize their responses to familiar stimuli by reducing redundant processing. However, repetition suppression effects are typically observed within a few seconds of a stimulus being repeated in early visual cortex, but can be considerably longer in other regions such as the orbitofrontal cortex (Barron et al., 2016). Though there have been reports of repetition effects spanning many days (van Turennout et al., 2000), we think it is unlikely that our observed phenomenon of increased variability in temporal dynamics is related to the same mechanism purported to underlie repetition suppression effects (Barron et al., 2016; Grill-Spector et al., 2006). An alternative possibility is that these effects may be more consistent with patterns observed in medial temporal lobe, in which activity appears to decrease with repeated presentations of a stimulus that may underlie a sense of familiarity (Henson et al., 2000; M. Suzuki et al., 2011). As stimuli become more familiar, participants may attend to different features that they may have not previously noticed, or may focus more on internal thought such as mind wandering. Consistent with this interpretation, we observed a decreased ability to predict activity in early visual cortex from low level features such as image brightness the more times participants viewed a stimulus and also a decreased anti-correlation with the medial prefrontal cortex.

In contrast, ventromedial prefrontal cortex (vmPFC) showed a more pronounced temporal proximity effect, where sessions closer in time exhibited more similar neural patterns regardless of the number of viewing times. This finding illuminates the dynamic nature of interpretive processing: the meaning we derive from sensory input depends more on our current context—our goals, mood, and recent experiences—than on mere familiarity with the stimulus. The vmPFC’s established role in value computation and affective meaning generation (Roy et al., 2012; Ashar et al., 2017) makes it particularly sensitive to these fluctuating internal states, likely facilitated by its afferent connections from heteromodal association cortex and thalamic and limbic regions (Mesulam, 1998; Morecraft et al., 1992). Consistent with this interpretation we observed that this change accompanied changes in participants preferences of the videos themselves. Participants ranked ordered preferences for the videos were more consistent with viewing sessions that were in close temporal proximity. This challenges the assumption that people have stable and consistent transitivity of preferences, which is at the heart of most decision theories such as expected utility theory (von Neumann & Morgenstern, 1944) and weak axiom of revealed preferences (Arrow, 1959; Samuelson, 1938).

These findings have broader implications for experimental designs. Any longitudinal study that plans to repeatedly present a passive viewing paradigm should not expect participants to treat each viewing session independently. It is an open question as to how much time would need to pass before these effects might be mitigated, if they are more pronounced with specific types of stimuli, or how much drifting attention, or other internal states might drive some of these effects. These types of questions would need to be addressed in future research. Studies investigating the impact of interventions that use the same stimuli should also include a temporal control.

In summary, we examined the impact of repeatedly viewing the same stimuli on neural dynamics. Our findings reveal that no two viewings are ever truly identical—each session is shaped by the brain’s dynamic processing. Sensory systems appear to adapt efficiently to repeated input, refining their responses, while higher-order interpretive systems continuously reshape meaning based on personal values and momentary states. We speculate that perception and understanding increasingly evolve over longer timescales (e.g., weeks, months, or years) suggesting that even when the external stimulus remains unchanged, the internal experience and personal meaning never stands still.

## Methods

### Participants

Twenty participants (mean age = 24.0 SD = 4.70; nine female) from the Dartmouth College community participated in the study for monetary compensation ($20/hour, extra bonus payment after completion of the 5th session was $150.00). All participants provided informed consent in accordance with the guidelines set by the Committee for the Protection of Human Subjects at Dartmouth College.

### Procedure

Participants underwent five fMRI scanning sessions (mean interval = 11.0 days, sd = 3.57). Each session consisted of five scanning runs, during which participants watched a set of 24 videos (mean length = 228.17 seconds, SD = 69.42). The order of runs and videos was fixed across all sessions for each participant. The videos were selected to elicit a variety of affective experiences, including food (e.g., making cookies, ramen, and salad), social interactions (e.g., family, romantic), sports (e.g., soccer, skiing, bmx, and skydiving), and dance (e.g., ballet, modern, and aerial gymnastics).

### Revealed Preference Task

After each scanning session, participants completed a revealed preference task. Participants were presented with pairs of screenshots on a computer from two different videos they had just watched in the scanner and were asked to select the screenshot corresponding to the video they preferred. Participants indicated their preference for each pair of videos (276 comparisons). For each session, we computed the total number of times each video was preferred over all videos and computed an ordinal ranking of the videos based on the participant’s preferences. We assessed the stability of video preferences by comparing the ranks of each over viewing sessions.

### fMRI Data Acquisition

Structural images were acquired using a T1-weighted, single-shot, high-resolution MPRAGE sequence with an in-plane GRAPPA acceleration factor of 2 and prescan normalization: TR/TE = 2300/2.32 ms, flip angle = 8°, resolution = 0.9 mm^3^ isotropic voxels, matrix size = 256 x 256, and FOV = 240 mm x 240 mm.

Functional BOLD images were acquired in an interleaved fashion using gradient-echo echo-planar imaging (EPI) with prescan normalization, fat suppression, and an in-plane GRAPPA acceleration factor of 2: TR/TE = 2000/25 ms, flip angle = 75°, resolution = 3 mm^3^ isotropic voxels, matrix size = 80 x 80, FOV = 240 mm x 240 mm, 40 axial slices with full brain coverage and no gap, anterior-posterior phase encoding. Each session included five functional runs (total TRs per session = 2966; TRs per run: 588, 613, 598, 588, 579).

### Preprocessing

All MRI data from all the scan sessions underwent preprocessing using a custom pipeline developed with Nipype and NIPY (Gorgolewski et al., 2011; Jarrod Millman & Brett, May-June 2007). This pipeline included several steps to ensure the quality and consistency of the imaging data. First, the pipeline trimmed the first five non steady-state TRs from each functional scan. Subsequently, each functional volume was realigned to the mean functional volume using a two-pass procedure implemented via FSL’s MCFLIRT. Brain extraction (skull stripping) was then performed on T1-weighted structural images using ANTS (advanced normalization tools). Following this, transformations were calculated for linearly coregistering realigned functional volumes to skull-stripped structural images, and nonlinearly normalizing skull-stripped structural images to the ICBM 152 2-mm MNI (Montreal Neurological Institute) template. These transformations were concatenated and applied in a single step using basis spline interpolation in ANTS. Additionally, data were spatially smoothed using a 6-mm full width at half maximum Gaussian kernel. The preprocessing pipeline can be found at https://github.com/cosanlab/cosanlab_preproc.

After basic preprocessing steps, further denoising was performed via univariate voxel-wise general linear model (GLM) outlined in the DartBrains course (L. J. Chang et al., 2020). This entailed residualizing the signal associated with various covariates of no interest, including mean, linear, and quadratic trends, mean activity from a predefined CSF probability mask (thresholded at 0.85), 24 head motion parameters (six demeaned realignment parameters, their squares, their derivatives, and their squared derivatives), global signal-intensity and frame difference spikes (outlier cutoff = 3) using the nltools toolbox (L. Chang et al., 2018) (https://github.com/cosanlab/nltools).

We used a k=50 parcellation scheme (http://neurovault.org/images/39711) for spatial feature selection, which simultaneously reduces the overall computational complexity while maintaining some degree of spatial specificity (Jolly & Chang, 2021). This parcellation was created by performing a whole-brain clustering of functional coactivation patterns across over 10,000 published studies available in the Neurosynth database (de la Vega et al., 2016).

### Data Analysis

#### Between-subject and within-subject similarity analysis

We used pairwise inter-subject correlation (ISC) analysis (Nastase et al., 2019) to assess temporal neural similarity across participants during video watching. ISC was calculated for each video and then averaged across videos. Between-subject correlations (BSC) were quantified as the mean of the lower triangle of the Fisher’s z-transformed pairwise correlation matrix of temporal neural signals between each participant within each ROI. For the first session BSC, we computed ISC on the first time participants viewed the videos. For the averaged inter-subject similarity, we calculated the BSC separately for each viewing session, and then averaged these values over all sessions. This procedure was performed separately for each parcel across the brain.

Within-subject correlations (WSC) were computed similarly by averaging the lower triangle of the Fisher z-transformed session-by-session correlation matrix of temporal neural signals within one ROI for each individual watching each video. We then averaged WSC values across participants and videos to obtain a single within-subject similarity value for each ROI. We used a linear contrast to test for decreasing BSC across sessions and performed inferences over videos. In addition, WSCs were compared to BSCs by testing for a significant contrast across videos.

#### Decomposing within-subject session variation

To characterize changes in brain activity across repeated movie viewings, we examined within-subject session-by-session representational similarity matrices. This involved calculating the similarity of temporal brain activity patterns within each ROI for each participant across the five viewing sessions. To formally quantify the observed patterns in the within-subject representational similarity matrices, we employed a distance regression approach (Parkinson et al., 2017; Xie et al., 2021). This method allowed us to decompose the observed similarity matrices into contributions from two distinct prototypical patterns: temporal proximity and viewing repetition. We modeled the temporal proximity effect using a Toeplitz matrix, where each descending diagonal decreased monotonically from 1 to 0, reflecting the assumption that sessions closer in time would exhibit more similar brain activity patterns due to autocorrelation. We modeled the viewing repetition effect using a matrix where values decreased monotonically from 1 to 0 as a function of distance from the first viewing session. This captures the idea that brain activity might change systematically with each repeated viewing, with the more viewings potentially becoming more distinct over time. For each participant and each video, we estimated the optimal linear combination of these two prototypical matrices that best reconstructed the observed within-subject representational similarity matrix. This was achieved by estimating β weights for each prototypical matrix using least squares minimization. β weights on the prototypical matrices reflect the degree to which the observed pattern of within-subject change over sessions could be independently explained by either the temporal proximity or viewing repetition effects. The contrast between the two effects reveals which region is better explained by one of the effects. We used a mixed-effects regression model to determine whether different brain regions exhibited different patterns of temporal proximity and viewing repetition effects. This model included random effects for both participants and videos to account for individual variability and video-specific effects. We specifically tested for an interaction between ROI (V1 vs. vmPFC) and effect type (temporal proximity vs. viewing repetition) to assess whether the influence of these effects differed between the two regions and extended this analysis across all 50 brain parcels.

#### Spatiotemporal analysis

Our primary analyses focused on the similarity of temporal dynamics averaged across voxels within each brain parcel. To investigate whether similar effects could be observed in spatial patterns, we extended our analysis to incorporate spatiotemporal dynamics (Chang et al., 2021). This involved creating for each ROI a session-by-session similarity matrix that captured the spatial similarity of voxel patterns separately for each timepoint in a video (i.e., TR). The correlation values of the lower triangle are converted to z scores using a Fisher r-to-z transformation and then averaged, and then inverted back from z-scores to r values. This provides a way to perform time-varying BSC when spatial similarity is computed across participants and WSC when spatial similarity is computed across viewing sessions for a single participant. Thus, these BSC and WSC metrics can vary over time for each video. The average over time within a movie and then across movies can also be computed to assess the total average spatiotemporal similarity. This approach complements the temporal similarity analyses by assessing if similar patterns are also present in the spatial patterns of voxels within a region.

#### Time interval sensitivity

To assess the robustness of our findings to variations in the time intervals between scanning sessions, we repeated our main analyses (within-subject session-by-session similarity and distance regression) using the actual time intervals between sessions for each participant, measured in days (mean interval = 11.0 days, SD = 3.57). This is in contrast to the primary analysis, which assumed equally spaced intervals.

Specifically, we first constructed a session-by-session time interval matrix for each participant, where each cell represented the number of days between a given pair of sessions. We then used these individual time interval matrices in two ways]. First, we directly correlated the session-by-session time interval similarity matrices (after inverting the distance matrices to reflect similarity) with the neural session-by-session similarity matrices for each ROI. This allowed us to assess whether the temporal proximity effect observed in our main analysis could be explained simply by the varying time intervals between sessions. Second, we incorporated the actual time intervals into our temporal proximity and viewing repetition models. Instead of assuming a linear decrease in similarity across equally spaced sessions, we used the time interval matrices to model the decay of similarity as a function of the actual time elapsed between sessions. For the temporal proximity model, values in the session-by-session similarity matrix decreased as a function of the time interval between those two sessions. For the viewing repetition model, values in the session-by-session similarity matrix decreased as a function of the time interval between each session and the first viewing session. We then re-estimated the β weights for these adjusted models using the same distance regression approach described previously. We used the same mixed effects regression approach as in our main analysis to test for differences in model parameters between V1 and vmPFC.

#### Assessing Changes in Perceptual Engagement

To investigate potential changes in perceptual engagement with repeated video viewings, we examined how brain activity tracked changes in video luminance across sessions. For each video, we calculated the average pixel intensity for each frame and downsampled this luminance time series to 0.5 Hz to match the sampling rate of our fMRI data. We then used a general linear model (GLM) to estimate the relationship between voxel-wise activity and the video luminance time series. This analysis was performed separately for each participant, video, and viewing session. To test for a systematic decrease in sensitivity to luminance across repeated viewings, we applied a linear contrast to the estimated GLM coefficients across sessions. Statistical significance was assessed using a one-sample t-test across participants, and the results were thresholded using a false discovery rate (FDR) correction of q < 0.05 to account for multiple comparisons.

#### Linking vmPFC Activity to Behavioral Preferences

To explore the relationship between neural activity and subjective preferences, we performed a representational similarity analysis (RSA) with video preferences estimated using the revealed preference task. For each session, we constructed a video-by-video revealed preference matrix, where each cell represented the number of times a participant preferred one video over another. This matrix was then used to derive an ordinal ranking of videos based on preference for each participant in each session. We then computed a session-by-session representational dissimilarity matrix for each video based on the pairwise distance using the Euclidean distance of the video ranks over sessions, and transformed the dissimilarity to similarity matrix.

To link neural activity to these preference patterns, we used RSA to compare the session-by-session preference similarity matrix to the session-by-session neural similarity matrix for each brain parcel (using the 50-parcel whole-brain parcellation described previously). This involved calculating the similarity using a Spearman correlation between the lower triangles of the two matrices. This analysis was performed separately for each participant. We specifically tested whether the ventromedial prefrontal cortex (vmPFC) exhibited a stronger relationship with preference changes compared to V1 using a paired-samples t-test. We also conducted a whole-brain analysis using the same RSA approach, applying a false discovery rate (FDR) correction of q < 0.05 to account for multiple comparisons across all 50 parcels.

## Acknowledgments

This work was funded by awards from the National Institute of Mental Health (R01MH116026 & R56MH080716). This work was also supported by funding from the National Science and Technology Council and National Taiwan University in Taiwan (NSTC 111-2423-H-002-008-MY4 and NTU 114L7870 to P.-H.A. Chen).

## Supplementary Figures

**Figure S1:**
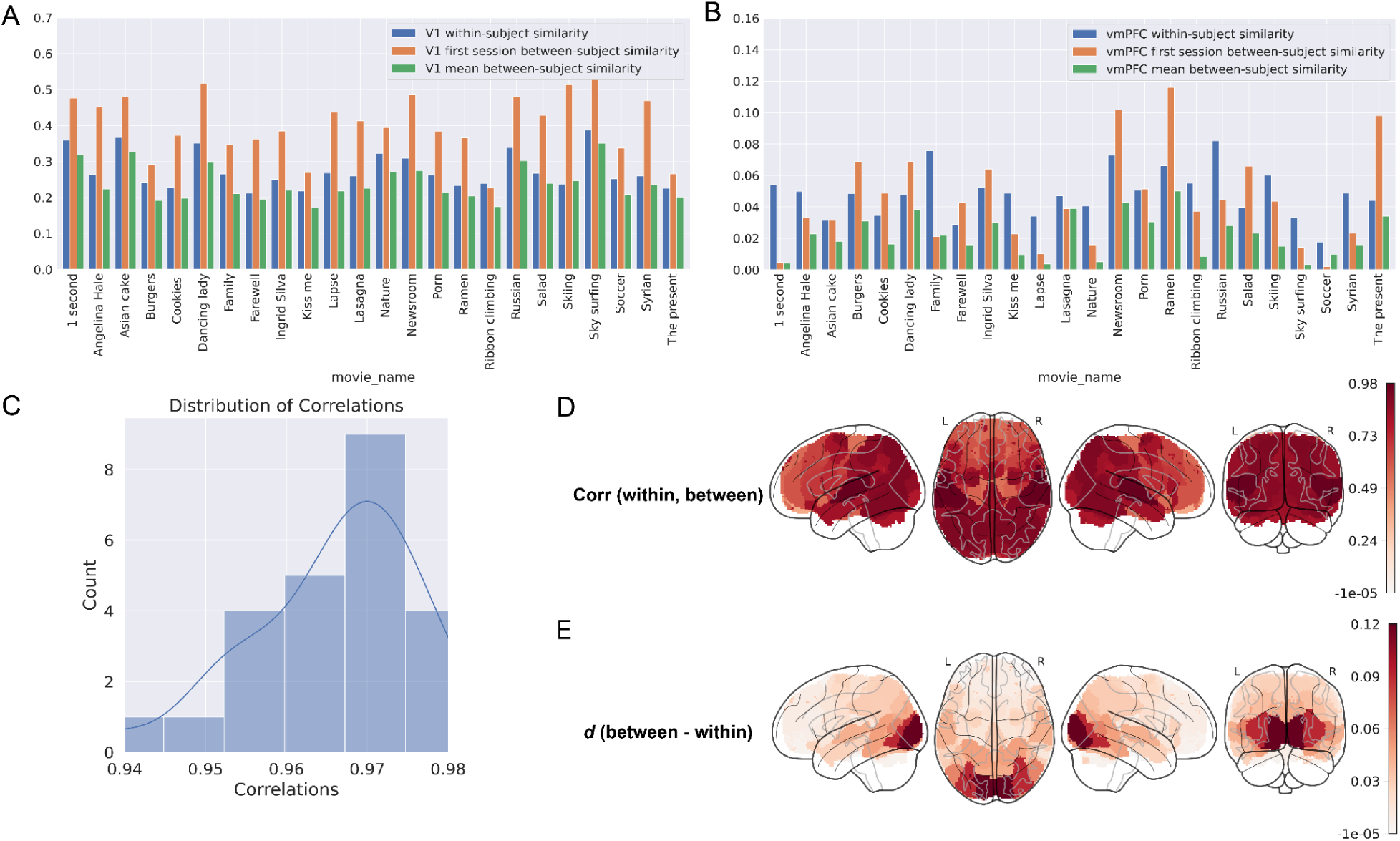
Comparison of within-subject and between-subject correlations across regions and videos. (A–B) Bar plots showing within-subject similarity (blue), first-session between-subject similarity (orange), and mean between-subject similarity across five sessions (green) for each video in V1 (A) and vmPFC (B). V1 exhibited more consistent WSC and BSC patterns across videos compared to vmPFC. (C) Distribution of spatial correlations (across parcels) between WSC and BSC maps computed separately for each video, showing overall high global correspondence. (D–E) Whole-brain maps showing the spatial correlation between WSC and first session BSC patterns across parcels (D) and the difference scores (first-session BSC minus WSC) across parcels (E).

**Figure S2:**
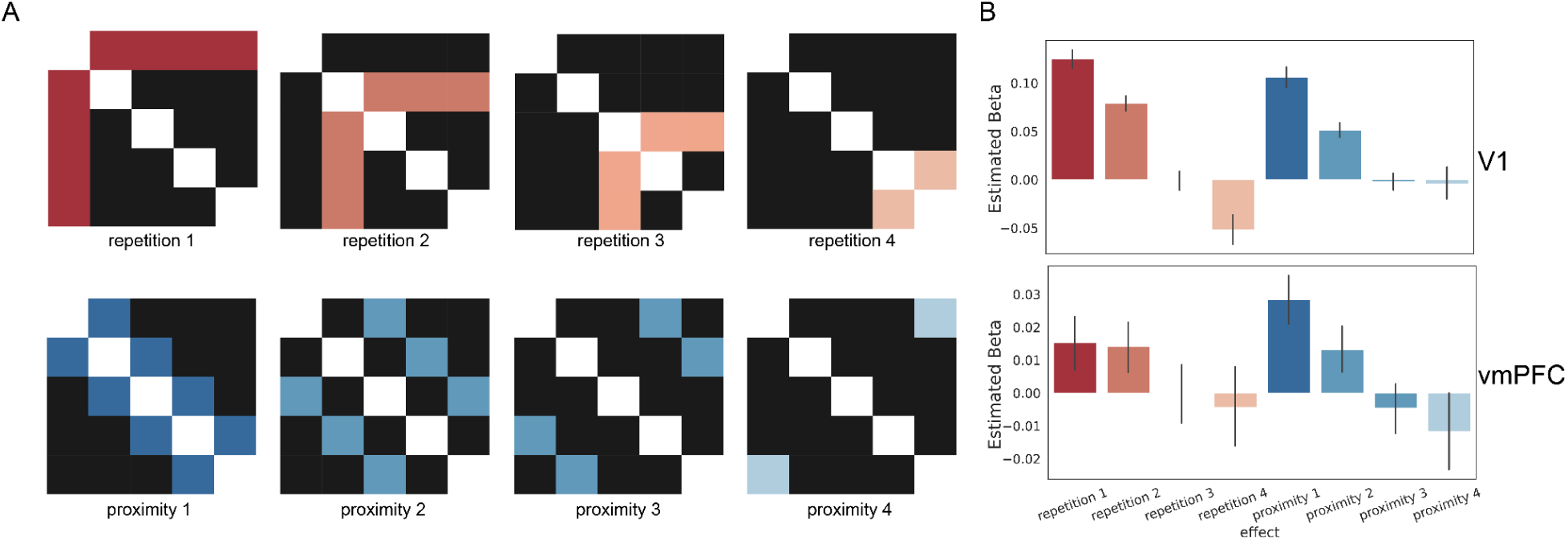
Individual viewing repetition and temporal proximity effects and their estimated contributions. (A) Prototypical session-by-session similarity matrices illustrating individual viewing repetition effects (repetition 1–4, top row) and temporal proximity effects (proximity 1–4, bottom row). Viewing repetition matrices model decreasing similarity as a function of distance from the first session, whereas temporal proximity matrices model decreasing similarity as a function of temporal distance between sessions. (B) Estimated β-weights for each repetition (1–4) and proximity (1–4) component from distance regression analyses in V1 (top) and vmPFC (bottom), showing their relative contributions to observed session-by-session similarity patterns across regions.

**Figure S3:**
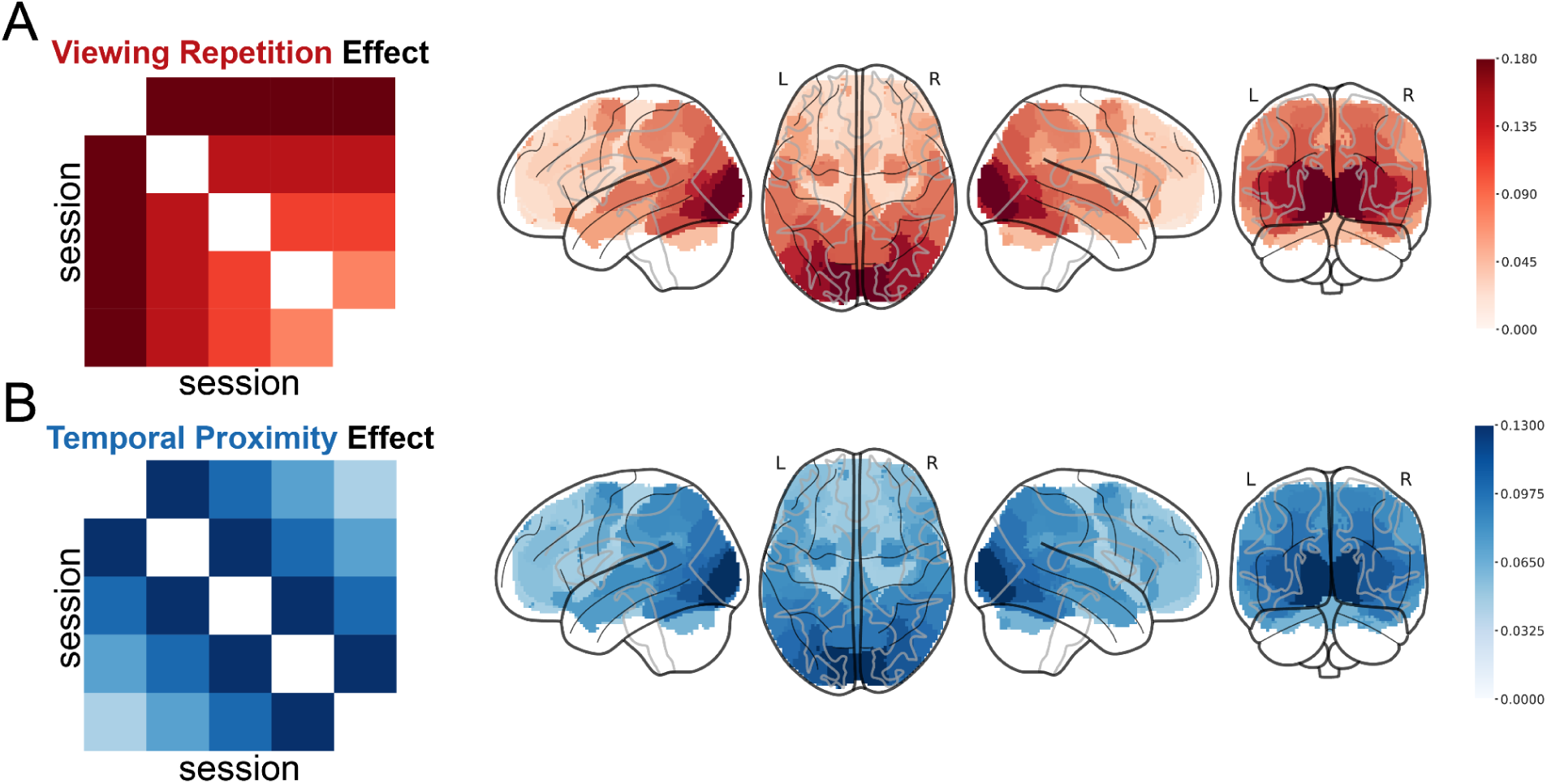
Global viewing repetition and temporal proximity effects across the brain. (A) Combined viewing repetition effect, and (B) Combined temporal proximity effect, showing the prototypical repetition and proximity models (left) and whole-brain maps of estimated β-weights (right).

**Figure S4:**
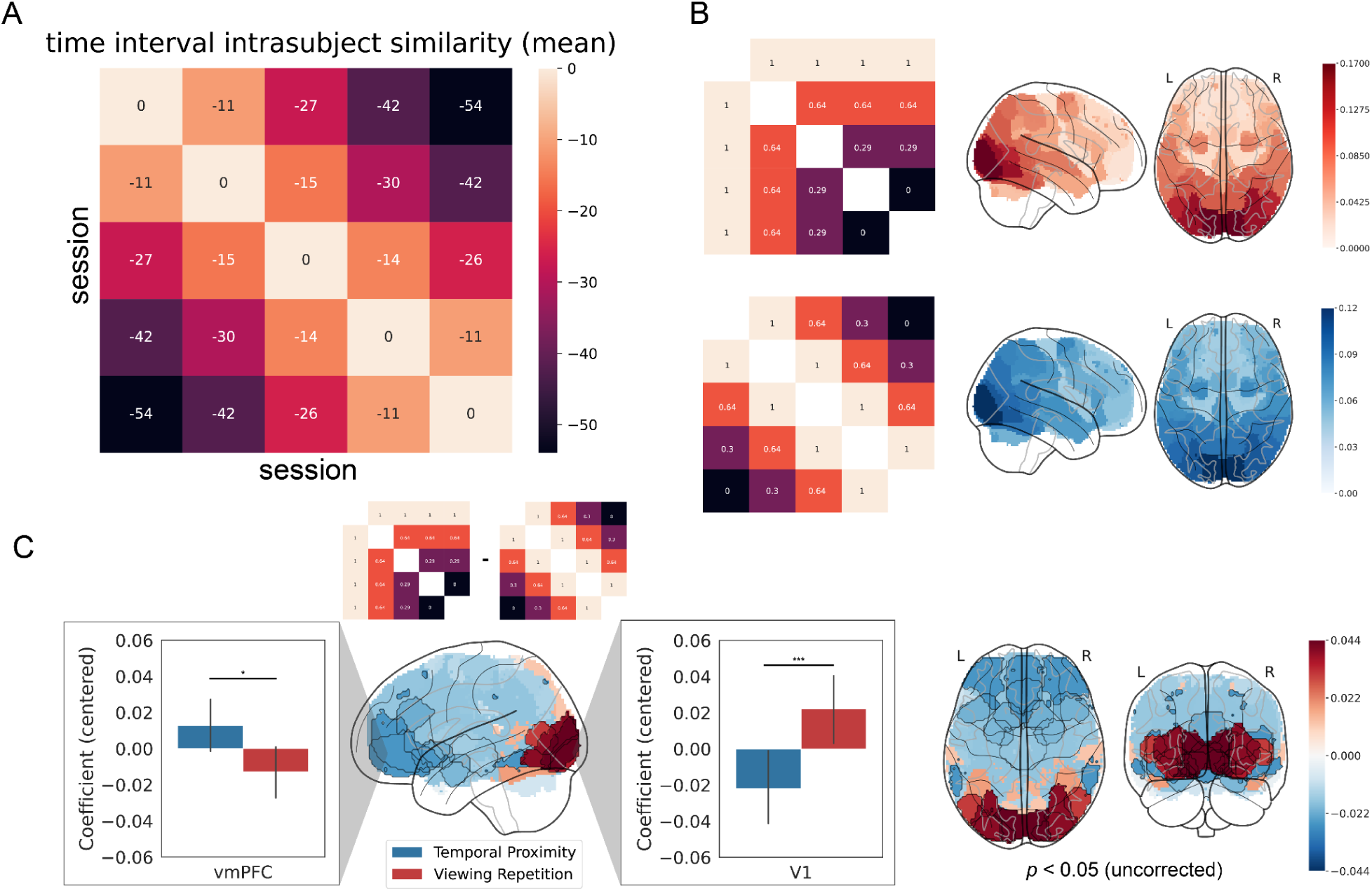
Time interval sensitivity analysis. (A) Average session-by-session time interval similarity matrix (in days) across participants. (B) Whole-brain analyses incorporating actual time intervals into the viewing repetition (top) and temporal proximity (bottom) models. Matrices on the left show the interval-weighted model structures, and brain maps on the right show the corresponding estimated weights across parcels. (C) Mixed-effects regression results comparing the contributions of interval-adjusted temporal proximity (blue) and viewing repetition (red) effects in V1 (right) and vmPFC (left), with whole-brain statistical maps showing regional distributions of these effects (*p* < 0.05, uncorrected). These analyses demonstrate that the main repetition and proximity effects remain robust when incorporating real temporal intervals.

**Figure S5.**
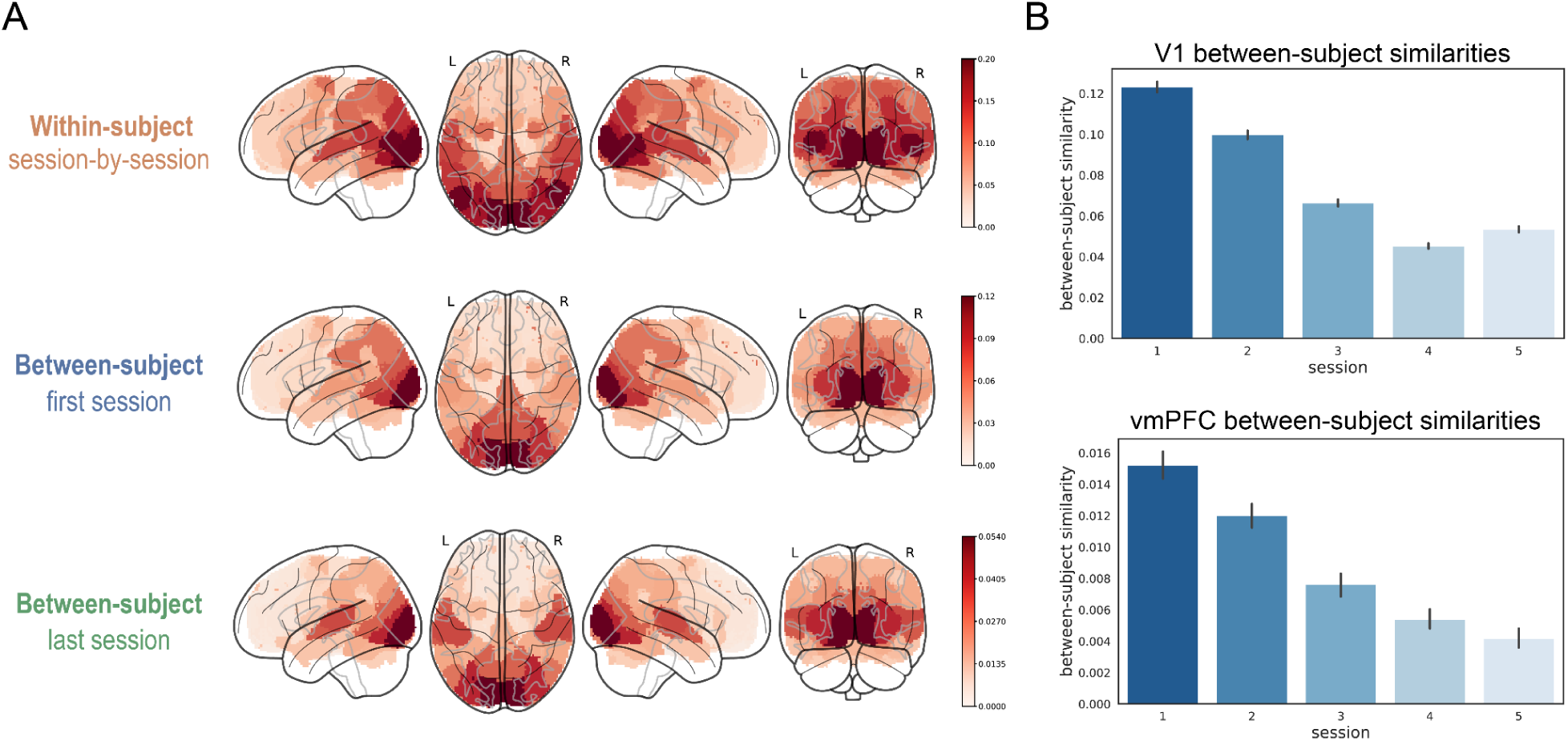
Spatiotemporal within- and between-subject correlation patterns across repeated viewings. (A) Whole-brain maps of within-subject similarity (top), first-session between-subject similarity (middle), and last-session between-subject similarity (bottom). (B) Session-by-session changes in between-subject spatiotemporal similarity in V1 (top) and vmPFC (bottom). Error bars represent bootstrapped 95% confidence intervals. These spatiotemporal analyses replicate the global WSC and BSC patterns observed in the main results, and show robust decreases in between-subject similarity across repeated viewings, particularly in V1 and vmPFC.

**Figure S6.**
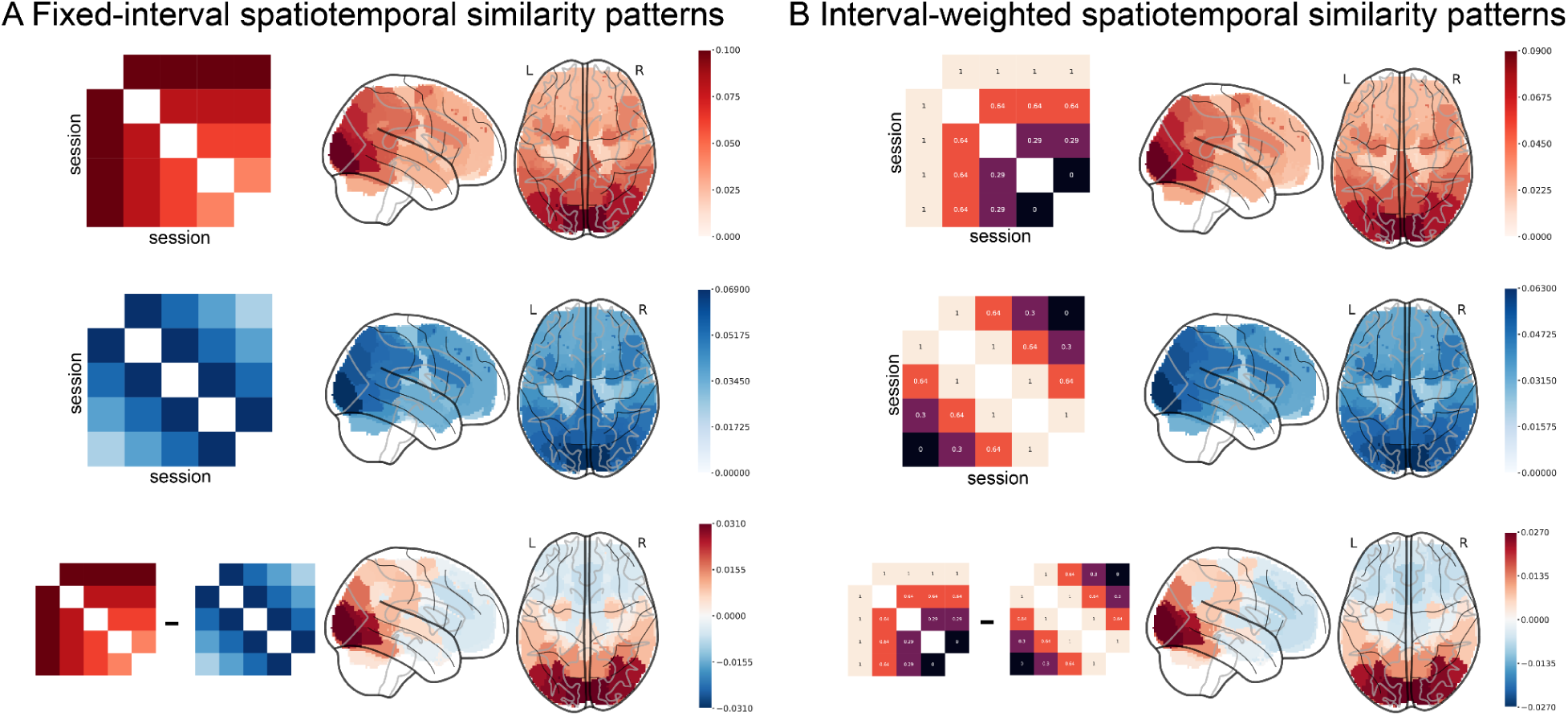
Replicating viewing repetition and temporal proximity effects using spatiotemporal analysis. (A) Results using spatiotemporal similarity matrices constructed with fixed, equally spaced session intervals. Left panels show the prototypical viewing repetition (top) and temporal proximity (middle) matrices and their difference (bottom), with corresponding whole-brain surface maps of estimated β-weights. (B) Results using spatiotemporal similarity matrices constructed with the actual temporal intervals (in days, normalized) between sessions. Left panels show the interval-weighted viewing repetition (top) and temporal proximity (middle) matrices and their difference (bottom), with corresponding whole-brain β-weight maps. These analyses replicate the results from the temporal analysis, showing that the repetition and proximity effects persist when assessed using spatiotemporal similarity patterns.

